# Multiple Displacement Amplification Facilitates SMRT Sequencing of Microscopic Animals and the Genome of the Gastrotrich *Lepidodermella squamata* (Dujardin, 1841)

**DOI:** 10.1101/2024.01.17.576123

**Authors:** Nickellaus G. Roberts, Michael J. Gilmore, Torsten H. Struck, Kevin M. Kocot

## Abstract

**Background:** Obtaining adequate DNA for long-read genome sequencing remains a roadblock to producing contiguous genomes from small-bodied organisms. Multiple displacement amplification (MDA) leverages Phi29 DNA polymerase to produce micrograms of DNA from picograms of input. Few genomes have been generated using this approach, due to concerns over biases in amplification related to GC and repeat content and chimera production. Here, we explored the utility of MDA for generating template DNA for PacBio HiFi sequencing using *Caenorhabditis elegans* (Nematoda) and *Lepidodermella squamata* (Gastrotricha).

**Results:** HiFi sequencing of libraries prepared from MDA DNA produced highly contiguous and complete genomes for both *C. elegans* (102 Mbp assembly; 336 contigs; N50 = 868 Kbp; L50 = 39; BUSCO_nematoda: S:92.2%, D:2.7%) and *L. squamata* (122 Mbp assembly; 157 contigs; N50 = 3.9 Mb; L50 = 13; BUSCO_metazoa: S: 78.0%, D: 2.8%). Amplified *C. elegans* reads mapped to the reference genome with a rate of 99.92% and coverage of 99.75% with just one read (of 708,811) inferred to be chimeric. Coverage uniformity was nearly identical for reads from MDA DNA and reads from pooled worm DNA when mapped to the reference genome. The genome of *Lepidodermella squamata*, the first of its phylum, was leveraged to infer the phylogenetic position of Gastrotricha, which has long been debated, as the sister taxon of Platyhelminthes.

**Conclusions:** This methodology will help generate contiguous genomes of microscopic taxa whose body size precludes standard long-read sequencing. *L. squamata* is an emerging model in evolutionary developmental biology and this genome will facilitate further work on this species.

## Background

In recent years, long-read technologies have become the industry standard for *de novo* genome sequencing. Pacific Biosciences’ (PacBio) multiple pass circular consensus sequencing (HiFi) has stood out as a popular choice for *de novo* genome sequencing due to its high accuracy (base-level resolution of >99%) and relatively long reads (∼10 Kbp) able to cover complex and repetitive genomic regions (Hon et al. 2020). Using adequate high-molecular-weight (HMW) DNA with as little damage as possible for sequencing library preparation is crucial to obtain reads that are both highly accurate and of adequate length (Pollard et al. 2018). Avoiding pooling multiple individuals serves to increase the assembly quality of genomes by reducing heterozygosity (Birky 1996; Flot et al. 2013; Huang et al. 2017; Simion et al. 2018). It follows that *de novo* genome projects are most likely to succeed when a large amount of unfragmented, HMW DNA with little to no exogenous contamination is available from a single individual.

Unfortunately, obtaining such amounts of starting material meeting the above requirements is not always possible. PacBio’s Ultra-Low DNA Input Workflow yields data volumes comparable to standard input libraries from as little as 5 ng of HMW template. This approach has successfully been used to produce high-quality genomes from individual small animals such as mosquitos (Kingan et al. 2019) and springtails (Schneider et al. 2021). However, even the Ultra-Low DNA Input Workflow is not applicable to very small animals (e.g., < 1 mm total length). For example, a single hermaphrodite specimen of *Caenorhabditis elegans* contains 959 somatic nuclei (Cohen-Fix and Askjaer 2017) and has a genome size of roughly 100 Mbp (*C. elegans* Sequencing Consortium 1998) meaning that a single specimen may contain as little as 200 picograms of DNA with comparable DNA content in other small-bodied phyla, also ranging in the hundreds of picograms of maximal yield. Thus, obtaining sufficient starting material meeting even the greatly reduced input requirements for the PacBio Ultra-Low DNA Input Workflow is not possible for many organisms. Further, if researchers want to avoid pooling multiple individuals or if the organism of interest is both small and rare, reduced input strategies are not an option thus far.

It is of little debate that for most applications leveraging genomes, assemblies that are both contiguous and well-annotated are ideal (reviewed by Laumer 2018). However, due to the challenges associated with sequencing genomes of small-bodied animals, metazoan genome sequencing efforts have been biased towards large-bodied organisms. Of the roughly 30 metazoan phyla, nearly all those that are still lacking high quality genome assemblies – or any genomic data at all in some cases – are small-bodied (Worsaae et al. 2023). For example, within Lophotrochozoa (=Spiralia) all phyla that are lacking in genomic data are mostly or exclusively small-bodied (Figure 1, Figure S1). With an average body size of <1 mm, these organisms are underrepresented in genomic and phylogenomic studies (Kocot et al. 2017; Laumer et al. 2019).

**Figure 1.**
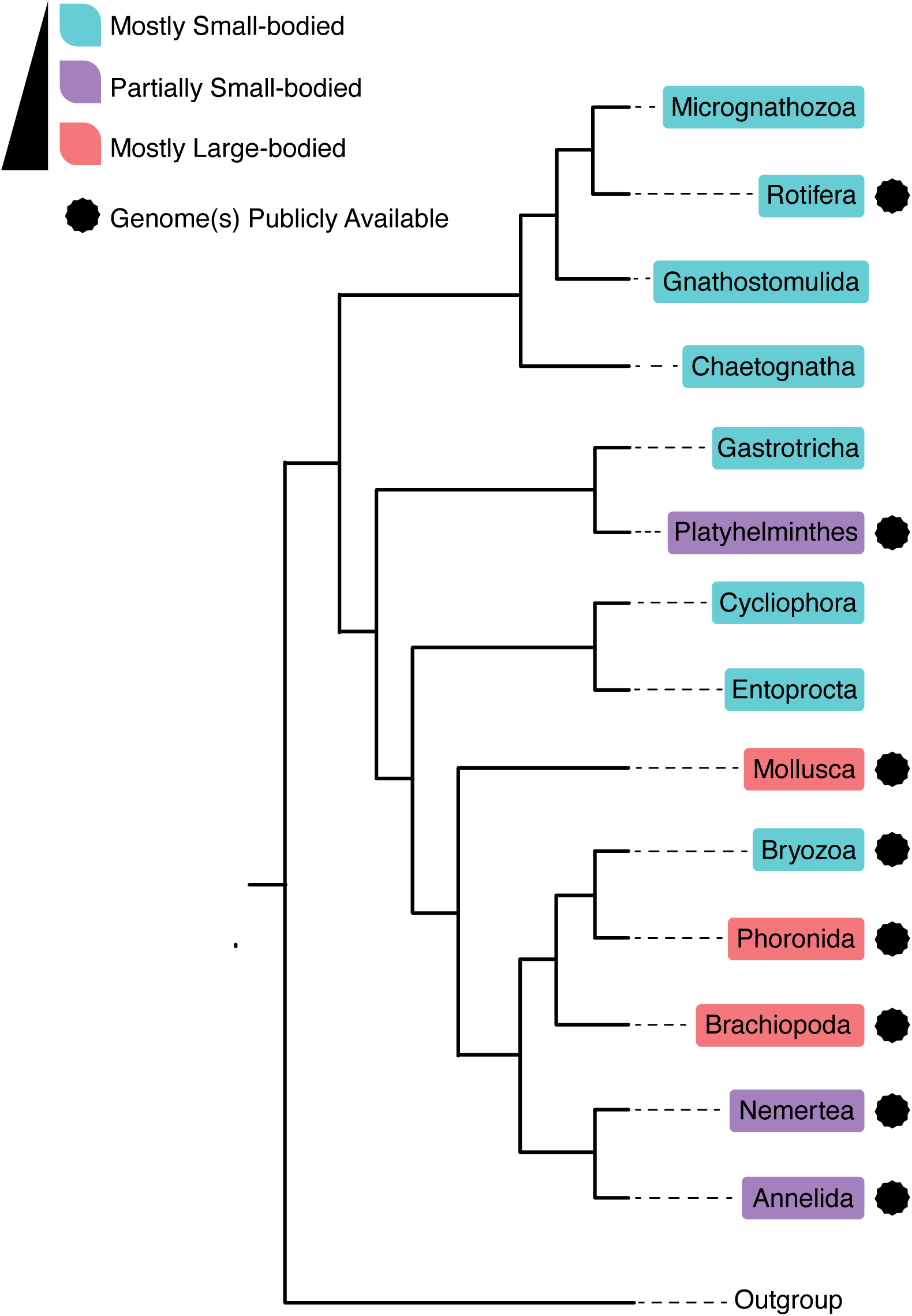
Phyla within Lophotrochozoa that are exclusively, partially, or mostly small-bodied. Those with one or more publicly available genomes are indicated. Tree modified from Laumer et al. (2019).

Whole genome amplification (WGA) methodologies are of great use to the fields of diagnostic medicine and single cell research (Evrony et al. 2015; Fu et al. 2015; Li et al. 2018; Zhou et al. 2020), but their utility in amplifying DNA for sequencing very small-bodied organisms in the age of long-read sequencing technologies has received little attention (but see Lee et al. 2023). WGA is a popular strategy for generating large amounts of DNA from limiting sample material through techniques such as multiple displacement amplification (MDA) (Dean et al. 2001), degenerate-oligonucleotide-primed-PCR (DOP-PCR) (Telenius et al. 1992), DOP-PCR (Telenius et al. 1992), PEP-PCR (Moghaddaszadeh-Ahrabi et al. 2012), LM-PCR (Carey MF et al. 2009), T-PCR (Tarr AW et al. 2005), and multiple annealing and looping based amplification cycles (MALBAC) (Chapman et al. 2015). However the use of WGA for long-read sequencing library preparation has largely been limited to the detection of structural variants, large deletions and inversions, and amplicon sequencing from degraded material (Evrony et al. 2015; Zhou et al. 2020).

WGA methodologies typically make use of PCR, isothermal amplification, or both (reviewed by Wang et al. 2022). PCR based protocols make use of specific primers, degenerate oligos, and/or repetitive genomic regions for priming. These strictly PCR-based amplification strategies are exponential and thus uneven or non-specific amplification may occur due to variation in template GC content and amplicon length. PCR-based methodologies have been used in medical and diagnostic approaches for their ability to analyze copy number variation and identify small structural variants or small stretches of SNP variation, but their utility for long-read sequencing is limited due to typical product fragment sizes only being within 100-1000 bp. Long-range PCR based approaches have been demonstrated to be successful for *de novo* genome sequencing of specimens in which material is limited (Laumer 2023; Stevens et al. 2023). Picogram input multimodal sequencing (PiMmS) makes use of oligo-dT beads to separate cDNA and genomic DNA from the same individual, providing transcript evidence from the same individual for downstream gene annotation. PiMmS amplification technique involves DNA purification using a salting out procedure to provide HMW ethanol-precipitated DNA. As PiMmS makes use of PCR, a qPCR step is required to estimate cycle number to reduce artifacts produced during PCR amplification.

Isothermal amplification in the case of MDA uses Phi29 DNA polymerase to amplify long fragments of linear or circular DNA (Dean et al. 2001; Dean et al. 2002; Lasken 2007). Amplification using Phi29 polymerase is highly accurate due to high strand-displacement activity, extremely high processivity, and strong proofreading activity (Garmendia et al. 1992). MDA, as opposed to PCR-based WGA approaches, has the benefit of producing highly accurate fragments with an average fragment length greater than 10 Kbp (up to 100 Kbp). Like other WGA approaches, MDA can produce tens of micrograms of DNA from picograms of starting material. MDA typically makes use of short, random exonuclease-resistant primers to allow the reaction to proceed isothermally. Like PCR-based methods, MDA is exponential leading to potential biases in coverage based on GC content (Lasken and Egholm 2003; Borgström et al. 2017). In GC-rich regions, there may be reduction of polymerase processivity leading to underrepresentation of that region during amplification. However, differential binding affinities of GC- and AT-rich primers can lead to GC-rich regions being preferentially amplified (Benita et al. 2003; Sahdev et al. 2007). Further, the denaturation strategy used prior to MDA (alkali, thermal or otherwise) may affect access and efficacy of DNA binding in GC rich regions. For example, KOH alkali denaturation has been shown to increase primer access to GC rich template DNA (Pinard et al. 2006). Such amplification bias with respect to GC richness is a function of reaction gain, defined as the ratio of DNA product mass over DNA template mass. Compared to MALBAC and PicoPLEX single cell, increased reaction gain in the case of MDA more negatively influences amplification coverage, uniformity, and the detection of copy number variants (CNVs). However, unlike both MALBAC and PicoPLEX, MDA reactions with high gain do not have a meaningful increase in the rate of single nucleotide errors (de Bourcy et al. 2014). Fortunately, lowering amplification time reduces reaction gain, thus reducing the over-amplification of certain genomic regions (Dean et al. 2001), making MDA protocols with more modest amplification time preferable to generate template for de novo sequencing. Lowering reaction gain also can also reduce chimera formation. During MDA, chimeric DNA fragments can form when 3’ termini are displaced and anneal to nearby displaced 5’ strands, forming chimeric DNA fragments that are eventually made into double stranded DNA after random hexamer annealing and extension (Lasken and Stockwell 2007). Previous studies have suggested a rate of up to 1 chimera/10 Kbp for MDA, which is problematic for the recovery of specific genomic regions, the analysis of copy number variation and *de novo* assembly, although these problems can be reduced with adequate coverage (Rodrigue et al. 2009). Proper preservation of samples and HMW DNA extraction (resulting in fewer 3’ termini) can reduce chimera formation (Nelson 2014), especially when combined with lower reaction gain. Moreover, as MDA can be applied directly to lysed cells (Hosono et al. 2003), shearing of DNA during extraction can be avoided altogether when aliquots of cell cultures or entire microscopic organisms are lysed followed by immediate amplification in the same tube. Taken together, MDA’s isothermal nature, high accuracy, production of fragments longer than typical PacBio HiFi reads, and ability to amplify DNA directly from freshly lysed cells makes it an attractive choice for template preparation for HiFi sequencing. Under low reaction gain and in low volumes, sequencing of MDA DNA can achieve high data uniformity and coverage (Li et al. 2018). Thus, MDA was chosen for this study.

Here, we investigated the utility of MDA for generating adequate template DNA for PacBio long-read sequencing from very small animals (<1 mm). We demonstrate the utility of this approach using the model organism *Caenorhabditis elegans,* which was selected because a high-quality reference genome and HiFi reads generated from unamplified DNA extracted from a pool of worms are already available for comparison. We additionally chose the gastrotrich *Lepidodermella squamata* (Dujardin 1841) as a test of MDA’s ability to assist in the *de novo* long-read sequencing of a non-model organism. Gastrotricha (Metschnikoff, 1865), colloquially referred to as “hairy bellied worms” is a phylum of approximately 749 species of microscopic benthic invertebrates found in aquatic habitats worldwide (Margulis and Schwartz 1998). Of these, 346 are found in freshwater habitats, with the rest inhabiting marine and brackish water. Gastrotricha is composed of two orders, Chaetonotida (Remane, 1925), which include both fresh and marine species, and Macrodasyida (Remane, 1925), which is an almost exclusively marine clade. Living mostly in sandy, benthic habitats, key taxonomic characters of Gastrotricha include the variable presence of scales, spines, or hooks on the body wall and the presence of adhesive tubes for attaching to substrate. Gastrotrichs typically range from 70 µm to 1 mm in size, although they may reach up to 3 mm in the case of the genus *Megadasys* (Schmidt, 1974, Fontaneto et al. 2015). Additionally, gastrotrichs are flattened along their dorsal-ventral axis reducing their volume even further in comparison to similar sized animals with a round body plan. The ∼250 µm freshwater gastrotrich *L. squamata* has proven to be a useful species for the study of Gastrotricha due to its ease of culture in a freshwater solution containing wheat grains inoculated with fungi, protozoa, and bacteria (Bennett 1979). Gastrotricha is of great interest to evolutionary biologists as its phylogenetic position has long been debated (Edgecombe et al. 2011; Struck et al. 2014; Laumer et al. 2015; Kocot 2016; Bleidorn 2019). Availability of a high-quality reference genome for *L. squamata* would also to the field of evolutionary developmental biology as *L. squamata* is an emerging model species (Hejnol 2015).

## Results

### Genome Assembly

Sequencing of a HiFi library generated from MDA DNA for *C. elegans* resulted in 708,811 HiFi reads (96x coverage of the 102.5 Mbp genome) with an average HiFi read length of 13,885 bp. The *de novo C. elegans* assembly was 102 Mbp contained in 336 contigs with an N50 of 868 Kbp and an L50 of 39. The *de novo* assembly was screened for contamination with Blobtools (Laetsch and Blaxter 2017), however all contigs had hits only to Metazoa (336) and thus none were removed. The *de novo* assembly had a BUSCO metazoa_odb10 score of 69.9% (Single copy BUSCOs [S]:67.5%, Duplicate BUSCOs [D]:2.4%) and a BUSCO nematoda_odb10 score of 94.9% (S:92.2%, D:2.7%) compared to 51.3% (S:50.7%, D:0.6%) and 98.5% (S:98.0%, D:0.5%), respectively, for the reference. The reference-guided *C. elegans* assembly was 102.5 Mbp contained in 281 contigs with an N50 of 16.9 Mbp and an L50 of 3 (Table 1).

**Table 1.** Assembly Statistics for *C. elegans* and *L. squamata*.

| *Organism:* | *HiFi* | *Genome* | *Contigs:* | *N50* | *L50* | *BUSCO* | *BUSCO* |
| --- | --- | --- | --- | --- | --- | --- | --- |
|  | *Reads:* | *Size* |  | *(Mb)* | : | *metazoa odb* | *nematoda odb* |
|  |  | *(Mb):* |  | : |  | *10* | *10* |
| *Lepidodermell* | 917258 | 122 | 157 | 3.9 | 13 | 80.8% |  |
| *a squamata* |  |  |  |  |  | (S: 78.0%, | N/A |
| *[MDA]* |  |  |  |  |  | D: 2.8%) |  |
| *Caenorhabditi* | 708811 | 102 | 336 | 0.87 | 39 | 69.9% | 94.9% |
| *s elegans* |  |  |  |  |  | (S: 67.5%, | (S: 92.2%, D: |
| *[MDA]* |  |  |  |  |  | D: 2.4%) | 2.7%) |
| *Caenorhabditi* | N/A | 102.5 | 281 | 16.9 | 3 | 51.3% | 98.5% |
| *s elegans* |  |  |  |  |  | (S: 50.7%, | (S: 98.0%, D: |
| *[Reference* |  |  |  |  |  | D:0.6%) | 0.5%) |
| *Guided MDA]* |  |  |  |  |  |  |  |

Sequencing of *L. squamata* resulted in 917,258 HiFi reads with an average HiFi read length of 13,358 bp. The *L. squamata* assembly was 147 Mbp contained in 527 contigs with an N50 of 2.9 Mbp and an L50 of 13. After filtering with Blobtools, 370 contigs were removed based on best hits to non-metazoans. The resulting assembly was 122 Mbp contained in 157 contigs with an N50 of 3.9 Mbp and an L50 of 13. BUSCO analysis of the decontaminated assembly using the metazoa_odb10 dataset was 80.8% (Single copy BUSCOs [S]: 78.0%, Duplicate BUSCOs [D]: 2.8%; Table 1).

### Coverage and Amplification Bias Estimation

Due to MDA’s potential for amplification bias with respect to GC and repeat content (Lasken and Egholm 2003; Borgström et al. 2017), we investigated the relationship between GC and repeat content and read coverage using the model organism *C. elegans* and its reference genome. To understand the level as to which MDA may have resulted in over amplification of genomic regions with respect to GC content, we compared mean coverage in 100 Kbp blocks of reads from our dataset based on MDA and reads from unamplified DNA extracted from a pool of worms (NCBI SRR22507561) mapped to the reference genome (GCA_000002985.3) (Figure 2, Figure S2). Due to the potential for repetitive regions to have GC content far outside the mean, we also investigated coverage with respect to repeat percentage. The reference *C. elegans* Bristol N2 genome has a GC content of 35% (GCA_000002985.3) and, according to RepeatModeler, it has a repeat content of 18.40%. Mean coverage was normalized for each dataset by subtracting the minimum mean coverage from each datapoint (representing a 100 Kbp region) divided by the range of the dataset [(x - min(x))/range(x)), x = coverage of 100 Kbp region]. Resulting normalized mean coverage, repeat content, and GC content in 100 Kbp blocks across all six chromosomes were plotted to visualize potential over-amplification of genomic regions during MDA (Figure 3, Figure 4A-B, Figure S3, Figure S4). This revealed that while some genomic regions showed over-amplification, amplification across the chromosome was consistent enough to provide adequate coverage. Plotting normalized coverage against GC content revealed virtually no difference between data generated from amplified and unamplified DNA (Figure 2) and the distribution was very uniform with respect to GC content with a few outliers. Surprisingly, the reads generated from unamplified DNA were more skewed towards higher coverage for GC values above the mean.

**Figure 2.**
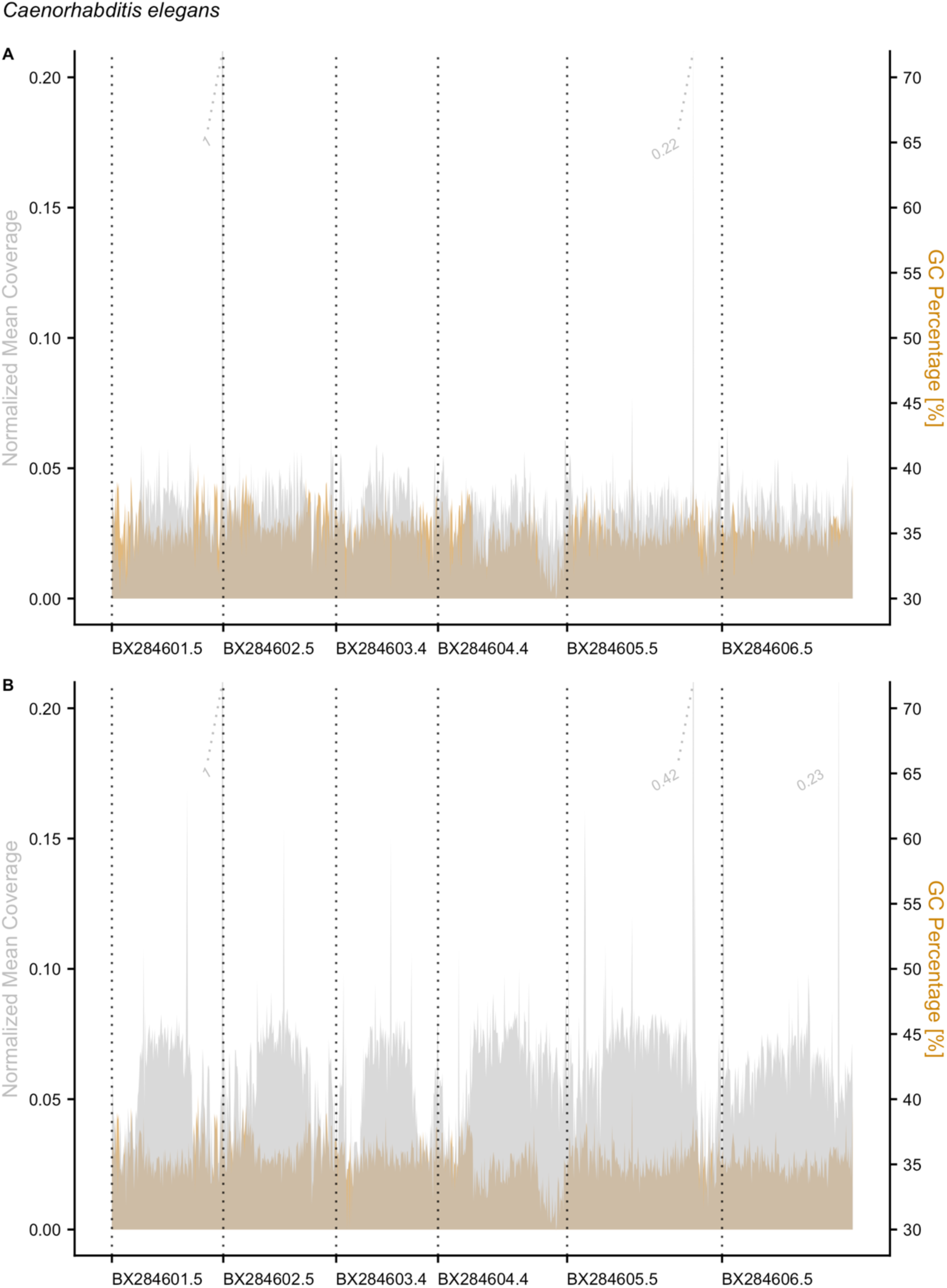
Mean coverage and GC content across 100 Kbp blocks of the *C. elegans* reference genome. A. Coverage of PacBio HiFi reads from MDA DNA aligned to the reference *C. elegans* genome [Gray] alongside GC percentage [Orange]. B. Coverage of unamplified PacBio HiFi reads aligned to the reference *C. elegans* genome [Gray] alongside GC percentage [Orange].

**Figure 3.**
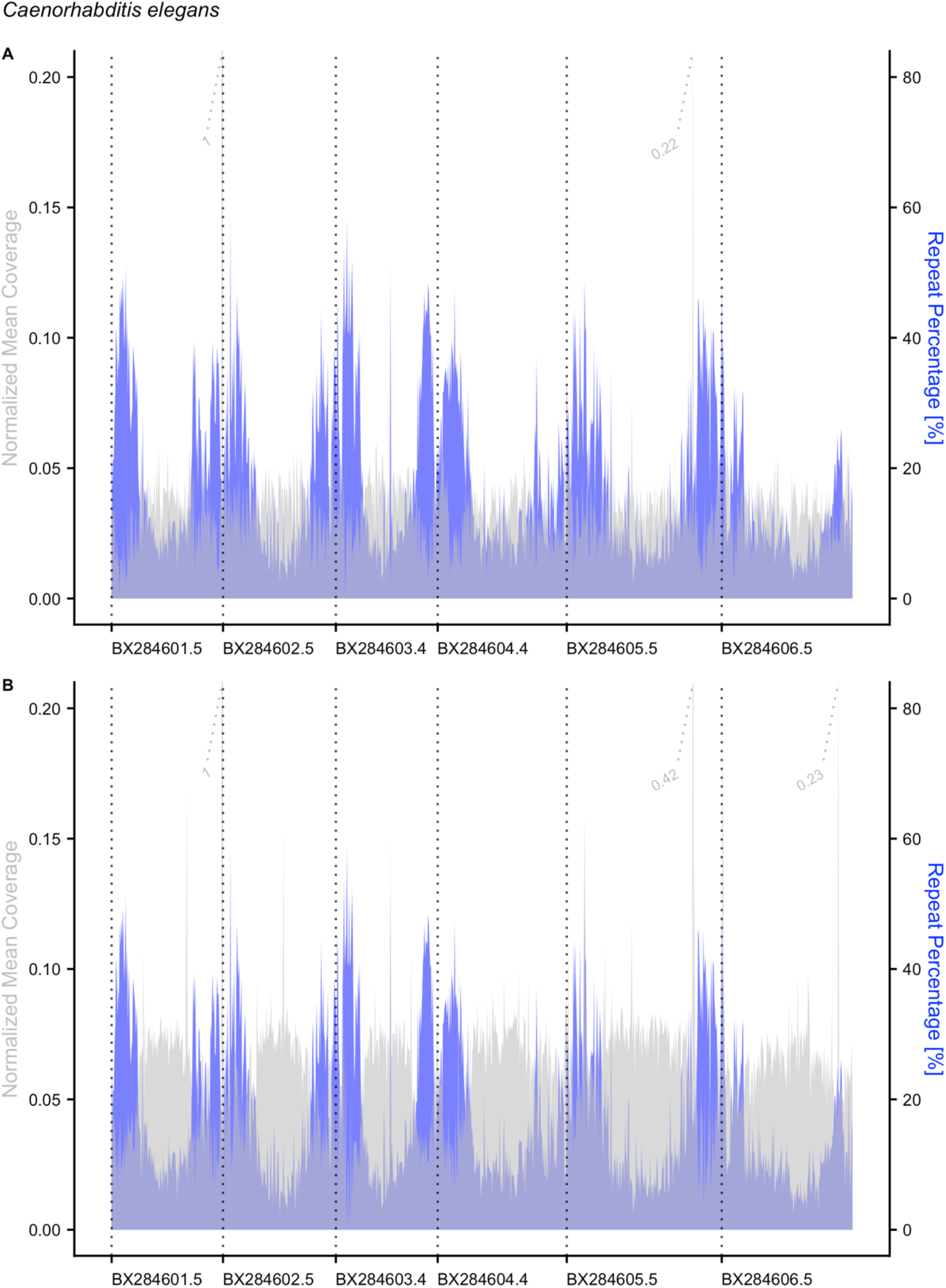
Mean coverage and repeat content across 100 Kbp blocks of the *C. elegans* reference genome. A. Coverage of PacBio HiFi reads from MDA DNA aligned to the reference *C. elegans* genome [Gray] alongside repeat content [Blue]. B. Coverage PacBio HiFi reads from unamplified DNA from a pool of worms aligned to the reference genome [Gray] alongside repeat content [Blue].

**Figure 4.**
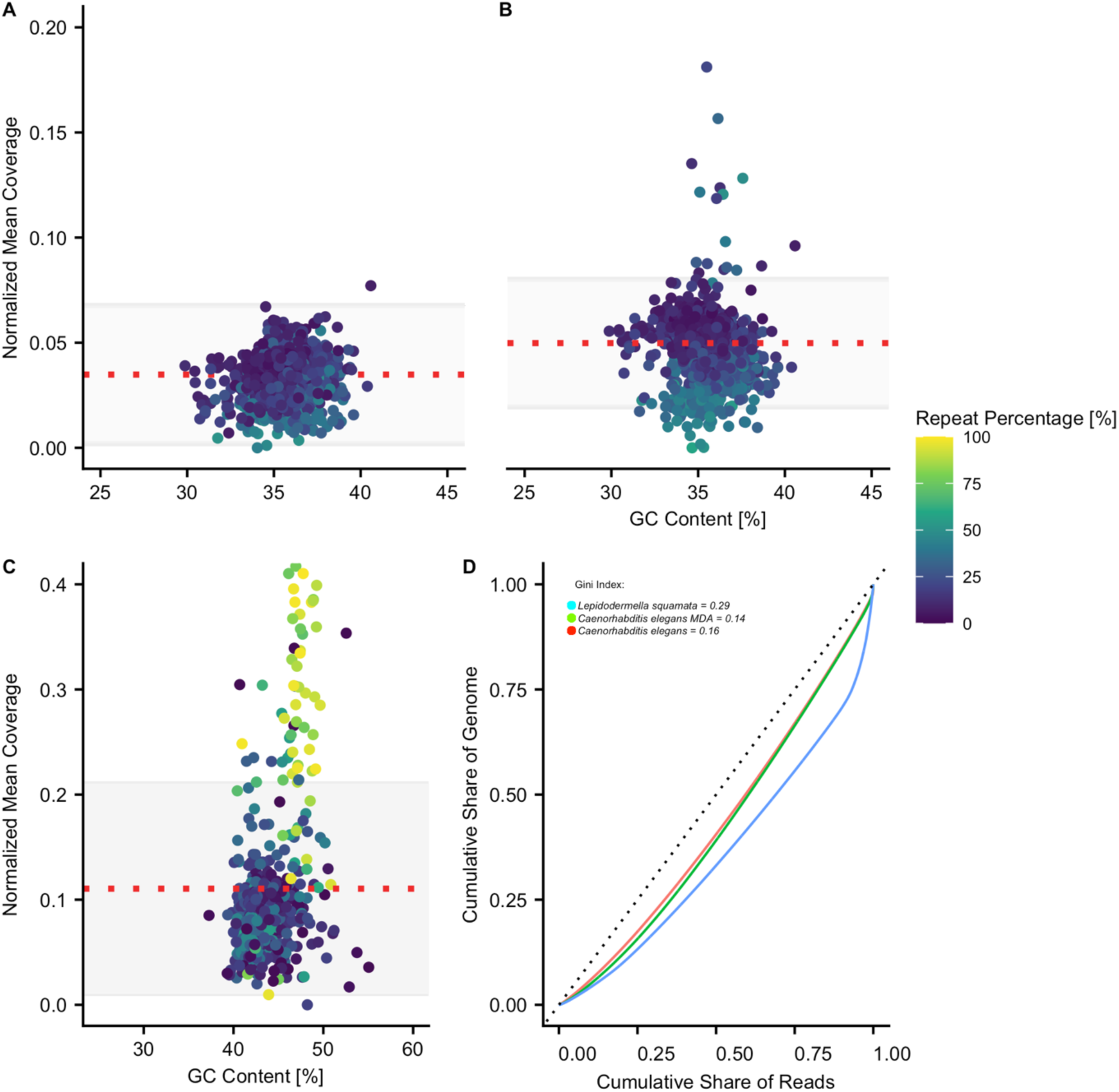
Normalized coverage, GC content, and repeat percentage of each genome assembly in 100 Kbp blocks. A. Dotplot showing coverage of MDA amplified reads with respect to GC content for 100 Kbp blocks of the *C. elegans* reference genome. B. Dotplot showing coverage of non-amplified reads from a pool of worms with respect to GC content for 100 Kbp blocks of the *C. elegans* reference genome. C. Dotplot showing coverage with respect to GC content for contigs in the *L. squamata* genome assembly based on MDA DNA. A-C. Repeat content of each contig is indicated according to the key at the top right of the figure. D. Lorenz curves and Gini values for each assembly indicating coverage uniformity. The diagonal line represents perfect read coverage uniformity.

Lorenz curves, defined here as a plot of the cumulative share of reads against cumulative share of genomic positions covered by those reads, ordered from highest to lowest (Zong et al. 2012; de Bourcy et al. 2014) were calculated (Figure 4, D). The Gini index is calculated as the area between the calculated Lorenz curve, from the dataset, and a straight Lorenz curve, representing perfect uniform coverage. In our case, a Gini index of zero would indicate perfect uniform coverage and a Gini index of 1 would indicate maximally non-uniform coverage. The Gini index values for the amplified and unamplified datasets were nearly identical, 0.14 and 0.16, respectively, revealing that the uniformity of coverage between the unamplified and amplified *C. elegans* read sets is similar. This suggests little to no effect of MDA amplification bias on coverage uniformity in the case of *C. elegans*. Coverage statistics, including mean coverage above and below mean GC content are summarized in Table 2.

**Table 2.** Coverage Statistics and Gini index values for C. elegans and L. squamata unmapped reads.

| *Mapped Reads:* | *Mean Coverage Across all 100 Kbp Blocks:* | *Mean Coverage of 100 Kbp regions above mean GC content.* | *Mean Coverage of 100 Kbp regions below mean GC content.* | *Gini Index :* |
| --- | --- | --- | --- | --- |
| *C. elegans*<br>[MDA] | 98 [SD +/-62] | 102 [SD +/-77] | 92 [SD +/-19] | 0.14 |
| *C. elegans*<br>[Unamplified] | 136[SD +/- 69] | 136 [SD +/- 77] | 136[SD +/- 19] | 0.16 |
| *L. squamata*<br>[MDA] | 117 [SD+/- 102] | 142 [SD +/- 135] | 118[SD +/- 42] | 0.29 |

Without a reference genome, no mapping comparisons could be made for *L. squamata*, however reads were mapped to the primary assembly after removal of contamination with Blobtools to assess coverage with respect to GC content and repeat content. The resulting genome assembly of *L. squamata* has a GC content of 43% and a repeat content of 21.98%. A Lorenz curve was plotted and the Gini index was calculated as described above. This resulted in a Gini index of 0.29 indicating less uniform coverage than both *C. elegans* reads based on MDA DNA and reads based on unamplified DNA (Figure 4D). *L. squamata* was comparatively not as uniform with respect to GC and repeat content as observed for *C. elegans* showing higher coverage in repetitive regions (Figure 4C), which also happen to have higher than mean GC richness. While few 100 Kbp blocks with average and lower GC content have higher fold coverage (Figure 5, Figure S5), a very substantial part of the coverage at GC content values above 45% exhibits very high coverage at a repeat content higher than 50% (Figure 6, Figure S6). This indicates possible preferential amplification of repetitive regions with high GC content. Coverage statistics, including mean coverage above and below mean GC content are summarized in Table 2.

**Figure 5.**
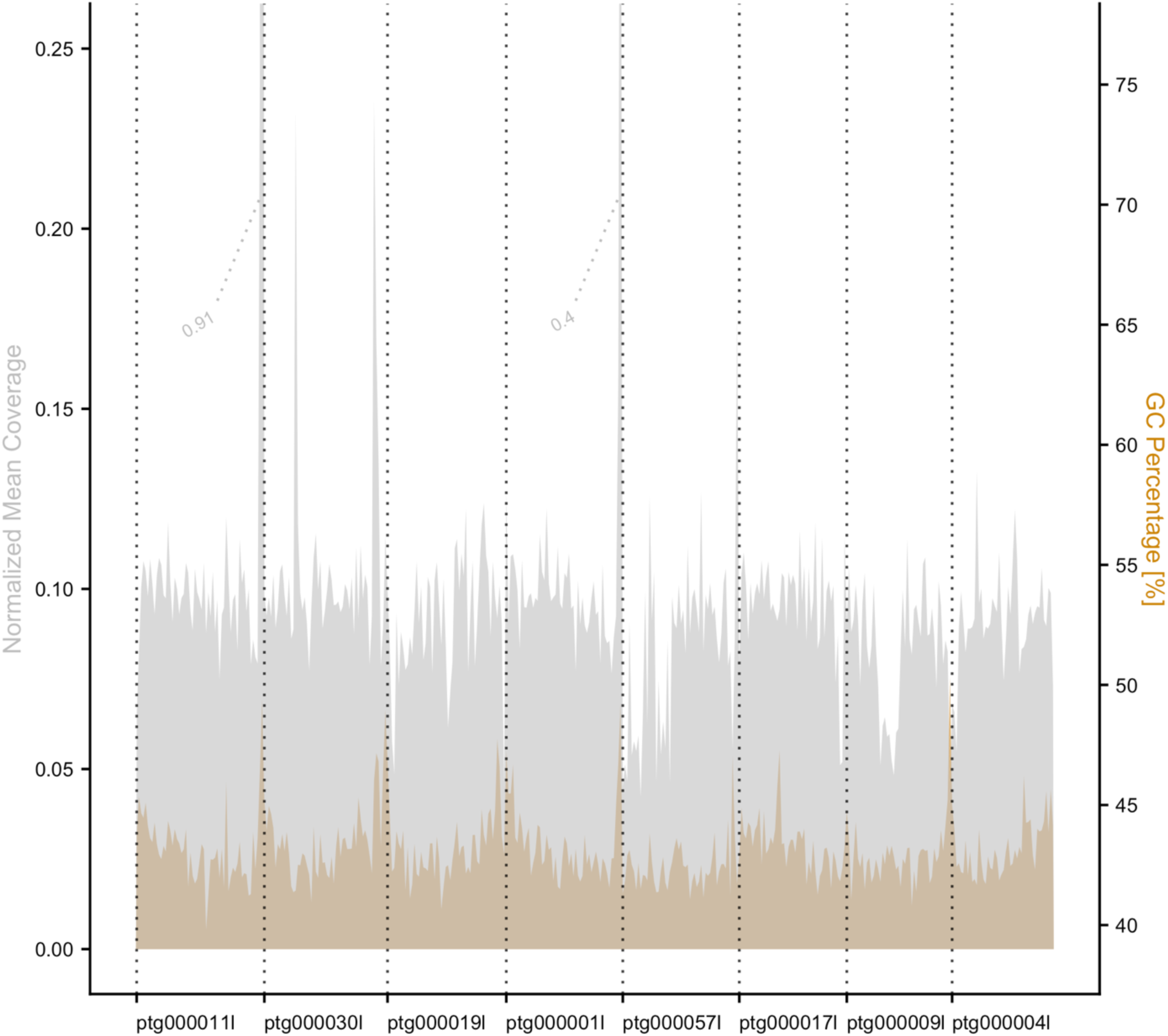
Coverage and GC content across 100 Kbp blocks of the eight largest contigs of the *L. squamata* genome. Coverage of PacBio HiFi reads from MDA DNA aligned to the 8 largest contigs of the *de novo L. squamata* genome assembly [Gray] alongside GC percentage [Orange].

**Figure 6.**
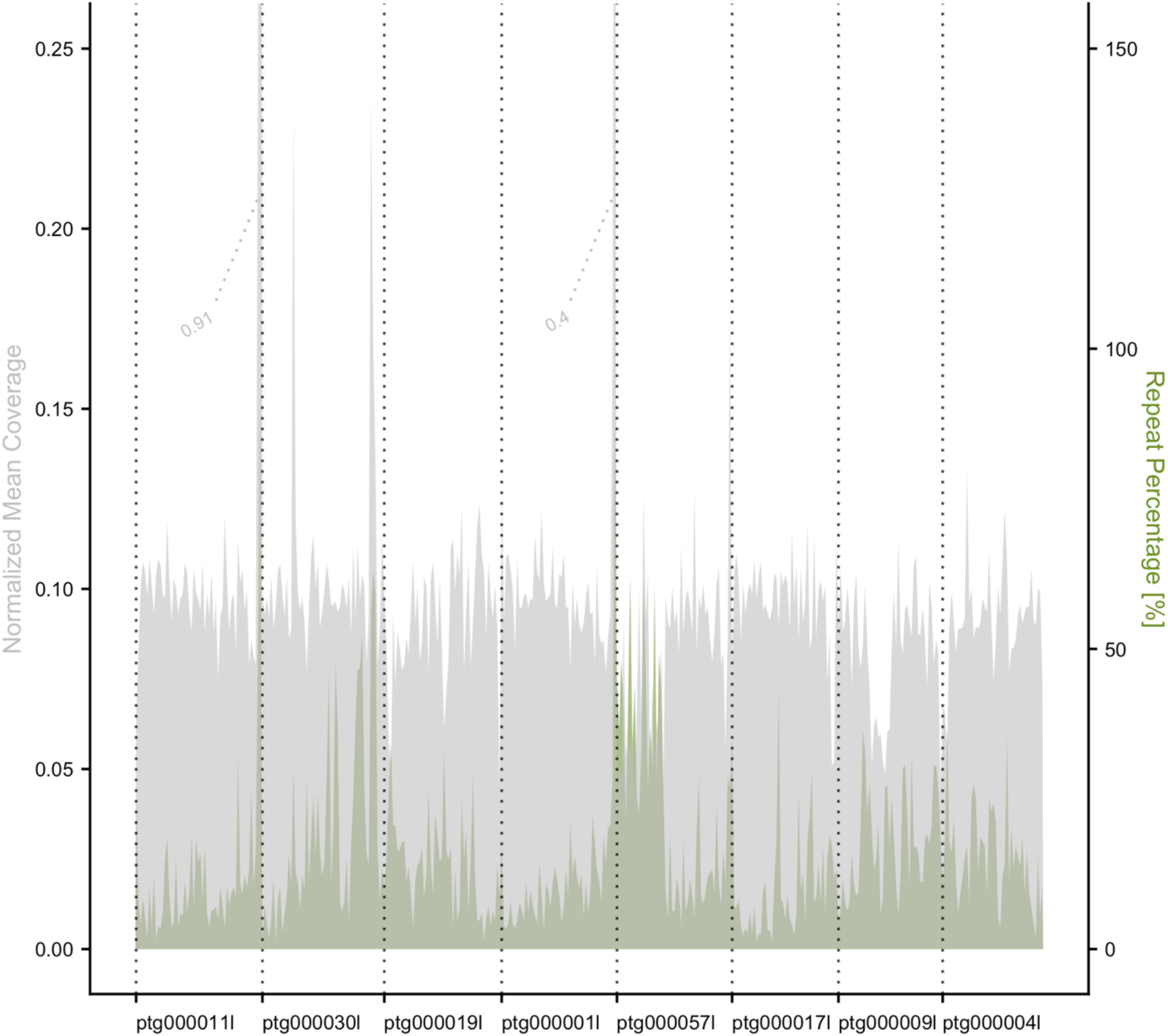
Coverage and repeat across 100 Kbp blocks of the eight largest contigs of the*L. squamata* genome. Coverage of PacBio HiFi reads from MDA DNA aligned to the 8 largest contigs of the *de novo L. squamata* genome assembly [Gray] alongside repeat percentage [Green]

### Annotation of *L. squamata* genome

BRAKER predicted 28,377 gene models with a BUSCO score of 79.80% (76.4% single-copy and 3.2% duplicated). The final assembly was estimated to be 21.98% repetitive. Of the 122 Mbp assembly, ∼16 Mbp (13.74%) was composed of unclassified repeats, while the remaining ∼26 Mbp (8.24%) was composed of simple repeats.

### Phylogenetic Analysis

To infer the phylogenetic position of Gastrotricha within Lophotrochozoa, we identified orthologs in the genomes of 23 lophotrochozoans (plus five outgroup taxa) broadly spanning the diversity of the group. We assembled a supermatrix of 2,779 genes totaling 676,632 amino acids with 18.68% missing data. Maximum likelihood analysis using the posterior mean site frequency model (PMSF) (Wang et al. 2018) fitting the C60 profile mixture model resulted in a tree with full bootstrap support for all nodes (Figure 7). Gastrotricha was recovered as the sister taxon of Platyhelminthes with maximal support, recapitulating the proposed sister-group relationship recovered previously (Struck et al. 2014) and dubbed Rouphozoa. Rotifera, the only representative of Gnathifera in this study, was recovered as the sister group to all other Lophotrochozoa with Rouphozoa as the next branching lineage and the sister taxon of all remaining taxa (Trochozoa) consistent with Struck et al. (2014). Within Trochozoa, we recover Mollusca as the sister taxon of a clade comprising, Annelida, Nemertea and the Lophophorate phyla (Brachiopoda, Bryozoa, and Phorinida), consistent with several previous studies (Struck et al. 2014; Kocot et al. 2017; Laumer et al. 2019). Due to the tree being based exclusively on genomes, several lineages of the superphylum, specifically most gnathiferan phyla (Chaetognatha, Gnathostomulida and Micrognathozoa), Cycliophora, and Entoprocta could not be included in this analysis.

**Figure 7.**
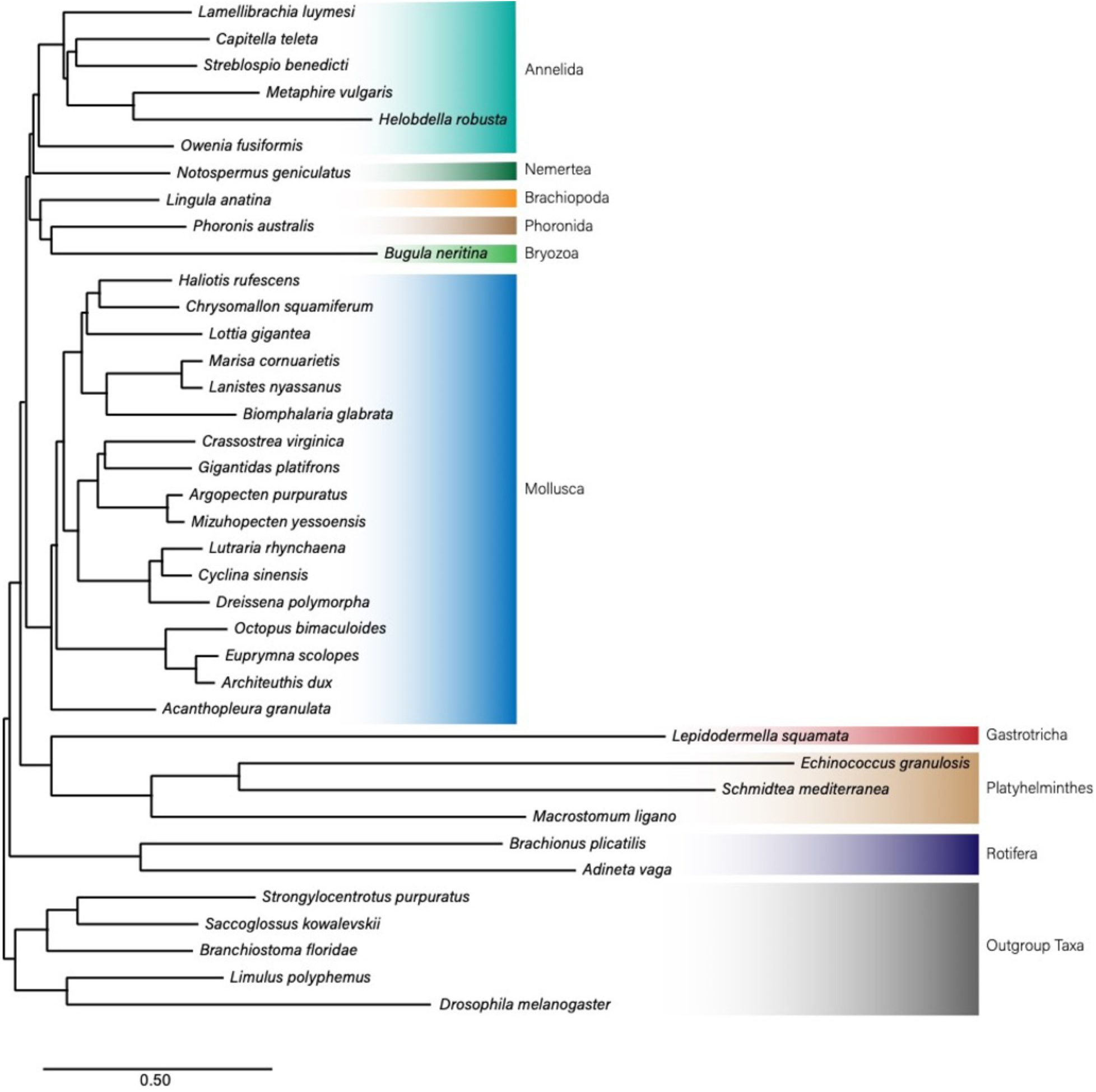
Evolutionary relationships of lophotrochozoan taxa with genomic data sequenced to date placing Gastrotricha as the sister taxon of Platyhelminthes (Rouphozoa). IQ-TREE 2 analysis of 676,632 amino acid positions with 18.68% missing data using the PMSF (LG+C60+F) with LG+C20+F as the guide tree. All nodes have 100% bootstrap support.

## Discussion

Here, we investigated the utility of MDA for generating adequate template DNA from small-bodied organisms for PacBio HiFi sequencing. Although coverage uniformity with respect to GC and repeat content are known biases of MDA, our results show that MDA of DNA obtained directly from lysed cells can be used to generate template for HiFi sequencing with limited coverage bias and artifacts.

Given existing high-quality genomic resources, *C. elegans* was chosen to explore the uniformity of coverage with respect to GC and repeat content by comparing our HiFi data generated from MDA DNA to HiFi data generated from unamplified DNA extracted from a pool of worms. Compared to HiFi reads generated unamplified DNA (Gini Index of 0.16), reads generated from MDA DNA (Gini Index of 0.14) performed similarly in terms of coverage uniformity, demonstrating that starting with samples as small as one-half *C. elegans* does not result in sequence data with significant amplification bias. While our *de novo C. elegans* assembly is not chromosome-level, it is rather contiguous and performed well in terms of BUSCO completeness (Table 1). When compared to PacBio Ultra-Low Input assemblies from single organisms, such as springtails (Schneider et al. 2021), our *C. elegans* assembly exhibits comparable completeness, albeit with lower contiguity. While our assembly performed well in terms of completeness (BUSCO Nematoda score of 94.9%), N50 was notably lower (868 Kbp) and the assembly consisted of a higher number of contigs (336) when compared to both springtail assemblies (N50 = 5.6 Mbp, 142 contigs, 211 Mbp size and N50 = 8.7 Mbp 165 Mbp size, 79 contigs for *Desoria tigrina* and *Sminthurides aquaticus* respectfully). This suggests that the PacBio Ultra-Low Input Workflow is likely a better choice than our MDA-based strategy for animals that can provide adequate HMW template DNA (>5 ng), but that would not be possible in the case of *C. elegans* and many (if not most) meiofaunal animals where a whole animal contains roughly an order of magnitude less DNA than required.

We were also concerned about the potential for formation of chimeras. However, investigation of inverted sequences and tandem regions with Alvis identified only one suspected chimeric contig in our *C. elegans* assembly and none in our *L. squamata* assembly, supporting previous hypotheses about chimera formation being highly correlated with high reaction gain (Lasken and Stockwell 2007; de Bourcy et al. 2014). Also, because we lysed cells and immediately proceeded with MDA in the same tube without a DNA extraction procedure, our template DNA underwent little handling that could cause breakage. This would result in fewer free 3’ ends of strands that could anneal to another strand and act as a primer to initiate chimera formation. As chimeric reads may be more problematic in data obtained from more fragmented template DNA, paired-end Illumina data generated from half of the specimen lysate before MDA could be used in future studies to aid in the detection of chimeras (see Lee et al. 2023). Examining the mapping of paired-end short reads to post-amplification HiFi reads could be used to reveal chimeric reads as one of the reads in a pair would be expected to map on one side of the chimeric junction while its mate would be expected to map elsewhere.

Our results demonstrate that this approach can be successfully used on small-bodied organisms without a reference genome to produce assemblies that are both rather contiguous and complete. The *L. squamata* assembly was rather contiguous with an assembly consisting of 157 contigs and an N50 of 3.9 Mbp approaching the level of contiguity produced using the PacBio Ultra Low Input Workflow (Schneider et al. 2021). The inferred BUSCO completeness score of 80.8% for *L. squamata* based on the metazoa_odb10 set suggests that this genome is lacking nearly 20% of the nuclear protein-coding genes thought to be single-copy in all metazoans. However, *L. squamata* is a long-branched taxon in our phylogenomic analysis and Gastrotricha as a phylum has been shown to be long branched in previous phylogenomic studies (Struck et al. 2014; Kocot et al. 2017; Laumer et al. 2019; Marlétaz et al. 2019). Moreover, there are no gastrotrichs or early-branching flatworms in the metazoa_odb10 reference dataset (Manni et al. 2021). Like the reference genome of *C. elegans* (BUSCO metazoa_odb10 score of 51.3%), *L. squamata* may have performed poorly in this BUSCO analysis due to sequence divergence rather than absence of these genes, although loss of BUSCOs in compact invertebrate genomes has been reported previously (Cunha et al. 2023). Regardless, the score recovered here represents a relatively high level of completeness when compared to other sequenced members of the lophotrochozoan superphylum (Figure S1). Annotation of the genome resulted in 28,377 genes, which BUSCO found to be 79.8% complete, which is comparable to the BUSCO score for the genome assembly, suggesting that the predicted gene model set is of high quality. Taken together, these results demonstrate that MDA combined with highly accurate HiFi sequencing can result in genome assemblies of similar quality to those of other non-model invertebrates (Rayko et al. 2020).

Despite the high contiguity of the *L. squamata* assembly, more bias with respect to repeat content and GC richness was observed in these data than in those from *C. elegans*. When compared to the *C. elegans* reference genome (18.40% repeats, 35% GC content), *L. squamata* (21.98% repeats, 43% GC content) has a higher repeat content. As a result of increased repeat percentage and higher overall GC content, we observed increased coverage in regions with >45% GC content, particularly in 100 Kbp blocks containing more than 50% repeats and in relatively short contigs. Amplification bias in GC-rich regions is a known artifact of MDA (Dean et al. 2002; de Bourcy et al. 2014), and our results provide supporting evidence for this. However, provided ample coverage, we were still able to capture enough breadth of the genome to produce a relatively complete and contiguous assembly.

MDA has been explored in the context of Oxford Nanopore (ONT) sequencing to produce relatively contiguous genomes from single nematodes (Lee et al. 2023). For *C. elegans*, the authors generated an assembly consisting of 499 contigs with an N50 of 656.2 Kbp and a BUSCO completeness score of 97.6%, compared to 336 contigs with an N50 of 868.1 Kbp and a BUSCO score of 92.2% in the present study. However, applying this approach to 13 other non-model nematodes, they recovered genome assembles ranging from 136.6 Mbp to 738.8 Mbp in 1,557 to 32,219 contigs with N50 values of 26.3 Kbp to 441.0 Kbp. Drops in coverage were observed around highly repetitive regions but coverage with respect to GC content was not examined in detail. Notably, T7 endonuclease digestion was needed in the case of ONT sequencing, as MDA’s secondary branching products can clog sequencing pores, reducing yield. Due to the nature of library preparation following shearing and size selection, endonuclease digestion is not needed for HiFi sequencing, a benefit of the approach presented here.

Given the successful application of this method to the organisms studied here, an important direction for future work would be to apply this strategy to other small-bodied phyla that are currently lacking published genomes (Chaetognatha, Gnathostomulida, Micrognathozoa, Entoprocta and Cycliophora). Increased genomic sampling of small-bodied phyla will lead to an elevated understanding about the evolution of small-bodied organisms and will assist in uncovering potential signatures and commonalities underlying the genomic architecture of small-bodied animals. The genomic consequences of miniaturization are poorly understood given the dearth of genomes from microscopic animals. Work to date has revealed that while some small-bodied lineages exhibit drastic simplification of genome architecture and reduction in content with respect to their large-bodied relatives (Seo et al. 2001; Mikhailov et al. 2016), have taken a more conservative route in regards to genome compaction (Martín-Durán et al. 2021; reviewed by Worsaae et al. 2023). Increased genomic sampling broadly spanning the metazoan tree stands to provide insight into genomic signatures of miniaturization. While our results demonstrate the utility of an MDA-based genome sequencing strategy for small-bodied animals with small genomes (i.e., 102 Mbp and 122 Mbp), the efficacy of this approach for organisms that are small-bodied, but with larger genomes still needs rigorous examination.

Combination of this approach with other low input sequencing techniques, such as PiMmS (Laumer, 2022), may be a valuable strategy as these techniques are expected to differ in their biases with respect to amplification uniformity. In contrast to long-range PCR amplification used in PiMmS and the PacBio Ultra-Low Input Workflow, the isothermal amplification used in MDA eliminates PCR duplicates at the cost of reduced uniformity across GC rich regions, particularly those high in repeats (Figure 3, Figure 4A, B). Here, DNA was amplified directly from lysed cells, bypassing HMW DNA extraction that may result in DNA loss from exceptionally small specimens and fragmentation, which we view as a beneficial characteristic of this approach. Careful consideration of the potential amount of DNA in a sample will aid in decision making regarding amplification directly from lysed cells as performed here versus HMW DNA extraction. Library diversity and complexity are both important considerations following any amplification protocol, as low complexity libraries, resulting from degraded or too little template DNA, can lead to poor quality assemblies. Finally, MDA’s yield of large amounts of DNA from limiting starting material allows multiple sequencing libraries to be produced from a single round of amplification, making it possible to increase coverage across under-amplified genomic regions, such as those with lower GC content.

## Conclusion

The amount of starting DNA has long been a limiting factor in *de novo* genome projects and, despite recent advances, many organisms still contain too little DNA for genome sequencing using available long-read sequencing library preparation protocols. In this study, the efficacy of whole genome sequencing following multiple displacement amplification with phi29 DNA polymerase was assessed. We demonstrate that HiFi sequencing data produced from MDA DNA can successfully lead to a fairly contiguous and complete assembly for both model organisms such as *C. elegans* (Nematoda) and for non-model organisms such as *L. squamata* (Gastrotricha) with limited coverage bias and an extremely low incidence of chimeras. This methodology could be further expanded upon or combined with other scaffolding techniques or amplification strategies to increase the contiguity of genomes from amplified material. This strategy has the potential to greatly expand the availability of whole genome sequencing data from a wide variety of DNA-limited sources including but not limited to other microscopic metazoans.

## Materials and Methods

### Laboratory Methods

Live cultures of *C. elegans* (Bristol N2) and *L. squamata* were purchased from Carolina Biological Supply Co. and kept alive in the laboratory under manufacturer recommended conditions. Half of a single individual *C. elegans* hermaphrodite (∼500 µm) cut with a sterile razor blade and a single individual *L. squamata* (∼190 µm) were added to 4 µl of Qiagen REPLI-g Advanced Single Cell Storage Buffer using a sterilized Irwin Loop as input for amplification using the Qiagen REPLI-g Advanced Single Cell kit following the manufacturer’s protocol (i.e., lysis with SDS and denaturation at room temperature, incubation at 30°C for 2 hours, 60°C for 10 minutes, and 4°C until purification). Resulting amplified DNA, 14.92 µg from *L. squamata* and 8.74 µg from *C. elegans* were sent to Hudson Alpha Discovery for sequencing on a shared single PacBio Sequel II flow cell, resulting in ½ a flow cell each.

### Assembly

HiFi reads were extracted using the pbbioconda extracthifi package and each set of reads was assembled with Hifiasm v.0.15.2. A reference guided assembly, using the genome of *C. elegans* Bristol N2 was generated for *C. elegans* using RagTag v.2.1.0 (-scaffold) (Alonge et al. 2022) with default settings. Genome statistics were generated using QUAST v.5.0 (Gurevich et al. 2013), and completeness was measured with BUSCO 4.0.2 (Simão et al. 2015). [Table 1,2] Completeness of all assemblies was analyzed with the BUSCO metazoa_odb10 BUSCO set, and additionally, both the *de novo* and reference guided *C. elegans* assemblies were assessed with the nematoda_odb10 BUSCO set. HiFi reads were mapped back to their respective assemblies using Minimap2 v.2.22 (Li 2018). The *C. elegans* and *L. squamata* assemblies were searched against the UniProt reference proteomes database with Diamond 2.0.14 (Buchfink et al. 2015) using the following settings: diamond blastx --db ∼/databases/uniprot_taxonmap.dmnd --query assembly.bp.p_ctg.fasta --outfmt 6 qseqid staxids bitscore qseqid sseqid pident length mismatch gapopen qstart qend sstart send evalue bitscore --sensitive --max-target-seqs 1 --evalue 1e-25 -- threads 32. The output of these tools as well as BUSCO were used as input for Blobtools. Further, the *de novo C. elegans* assembly was aligned to the publicly available reference *C. elegans* N2 genome (GCA_000002985.3) to assess completeness and screen for the presence of chimeric reads as identified by Alvis v.1.0 (Martin and Leggett 2021), a program that identifies chimeras in the assembly based on mapping statistics from mapping raw reads to the assembled genome. All *de novo* assemblies (*C. elegans* and *L. squamata*) were screened for contamination using Blobtools2 v2.6.1 by removing all contigs that had top BLAST hits to taxa other than Metazoa or “no-hit” when searched against the NCBI non-redundant protein database.

### Annotation

The repeat content of the contaminant filtered *L. squamata* genome was assessed using RepeatModeler (Flynn et al. 2020) and soft masked using RepeatMasker (Smit et al. 2015). Raw transcript reads from the publicly available *L. squamata* transcriptome (SRR1273732) were downloaded, quality trimmed with TrimGalore v.0.6.10 from NCBI and mapped against the soft masked genome using STAR (Dobin et al. 2013). The soft masked genome and mapped transcripts were given to the BRAKER (Hoff et al. 2019) pipeline as input to produce gene annotations with the following command: braker.pl --useexisting --cores 12 --softmasking -- UTR=on --crf --makehub --gff3 --species=Lepidodermella_squamata --genome Lepidodermella_sp.asm.bp.p_ctg_filtered.fasta.masked --bam RNAseq.bam. Gene annotations were measured for completeness using the BUSCO metazoa_odb10 database.

### Read Coverage Estimation

Assemblies of all three data sources (*L. squamata* and amplified and non-amplified *C. elegans*) were used as subjects to map HiFi reads as a query to estimate genome coverage and coverage bias. Following read mapping with Minimap2 v.2.22, Bedtools v.2.30.0 (Quinlan 2014) was used to separate the genomes into independent 100 Kbp windows in which both overall read coverage, mean coverage, repeat content and GC content were calculated. Resulting graphs were constructed in the R computing environment (R Core Team 2022). Lorenz graphs were constructed utilizing the package *gglorenz*. Additionally, we assessed coverage uniformity of our *C. elegans* HiFi data mapped against the reference genome, as well as PacBio HiFi reads generated from unamplified DNA extracted for a pool of worms of the same strain (SRR22507561). Read depth, breadth, and mapping rate for *C. elegans* to the reference assembly were calculated using SAMtools.

### Orthology Inference and Supermatrix Construction

To investigate the phylogenetic position of Gastrotricha, orthology determination and supermatrix assembly were performed following a modification of the pipeline of Kocot et al. (2017b) implementing new versions of the programs used if applicable. Lophotrochozoan genomes (including *L. squamata*) broadly spanning the diversity of the group were selected along with five outgroup taxa resulting in 28 OTUs. Homologous sequences were identified using OrthoFinder v.2.4.0 (Emms and Kelly 2019) and sequences shorter than 100 amino acids were removed, keeping the longest sequence. Homogroups were retained if they included greater than 75% of the total of all taxa, with those with less being removed. Fasta files were aligned with MAFFT v.7.310 (Katoh 2002), suspected mistranslated regions removed with HmmCleaner (Di Franco et al. 2019), and regions high in gaps or ambiguity were trimmed with BMGE v.1.12.2 (Criscuolo and Gribaldo 2010). Maximum likelihood gene trees were generated using IQ-Tree 2 (Minh et al. 2020) with the best-fitting model of amino acid substitution for each gene. Subsequent fasta files and trees were used as input for PhyloPyPruner v.0.9.5 (https://pypi.org/project/phylopypruner) to retrieve a set of strict [1:1] orthologs with the settings phylopypruner --min-taxa 15 --min-support 0.9 --mask pdist --trim-lb 3 --trim-divergent 0.75 -- min-pdist 0.01 --prune LS. The –min-taxa flag guaranteed removal of pruned alignments that did not include at least 15 taxa from the original dataset.

### Maximum Likelihood Estimation

The final supermatrix from PhyloPyPruner of 676,632 amino acid positions was used for phylogeny reconstruction in IQ-Tree2. Using maximum likelihood with the simplest profile mixture model available, LG+C20+F, a guide tree for PMSF (Wang et al. 2018) was produced. The PMSF analysis fitted the C60 profile mixture model, LG+C60+F, to the data and reconstructed the phylogeny assessing nodal support with 1000 rapid bootstraps.

## Availability of data and materials

Genomic data generated for this study have been deposited in the NCBI SRA and Genome databases under the BioProject accession number PRJNA1063365. Scripts used for assembly, annotation, read coverage investigation, and visualization are available at https://github.com/ngroberts/MDA_Hifi_Roberts_2023.

## Authors’ contributions

Study design, NGR, KMK. Genome assembly, sequencing, and annotation, NGR, KMK, THS. Statistical analyses and data visualization, NGR, MJG, THS. Manuscript preparation, NGR, KMK.

