## Supplementary material for "Multiple Displacement Amplification Facilitates SMRT Sequencing of Microscopic Animals and the Genome of the Gastrotrich *Lepidodermella squamata* (Dujardin, 1841)": Figure S1

**Table of Contents:**

Figure S1 - BUSCO scores of publicly available lophotrochozoan genomes (Metazoa odb10).

Figure S2 - Mean coverage, and GC content across 100kb blocks of the *C. elegans* reference genome with outliers removed.

Figure S3 - Mean coverage and repeat content across 100kb blocks of the *C. elegans* reference genome with outliers removed.

Figure S4 - Coverage and GC content across 100kb blocks of the eight largest contigs of the

*L. squamata* genome with outliers removed.

Figure S5 – Coverage and repeat across 100kb blocks of the eight largest contigs of the

*L. squamata* genome with outliers removed.

Figure S6 - Normalized coverage, GC content, and repeat percentage of each genome assembly in 100kb blocks with outliers removed.

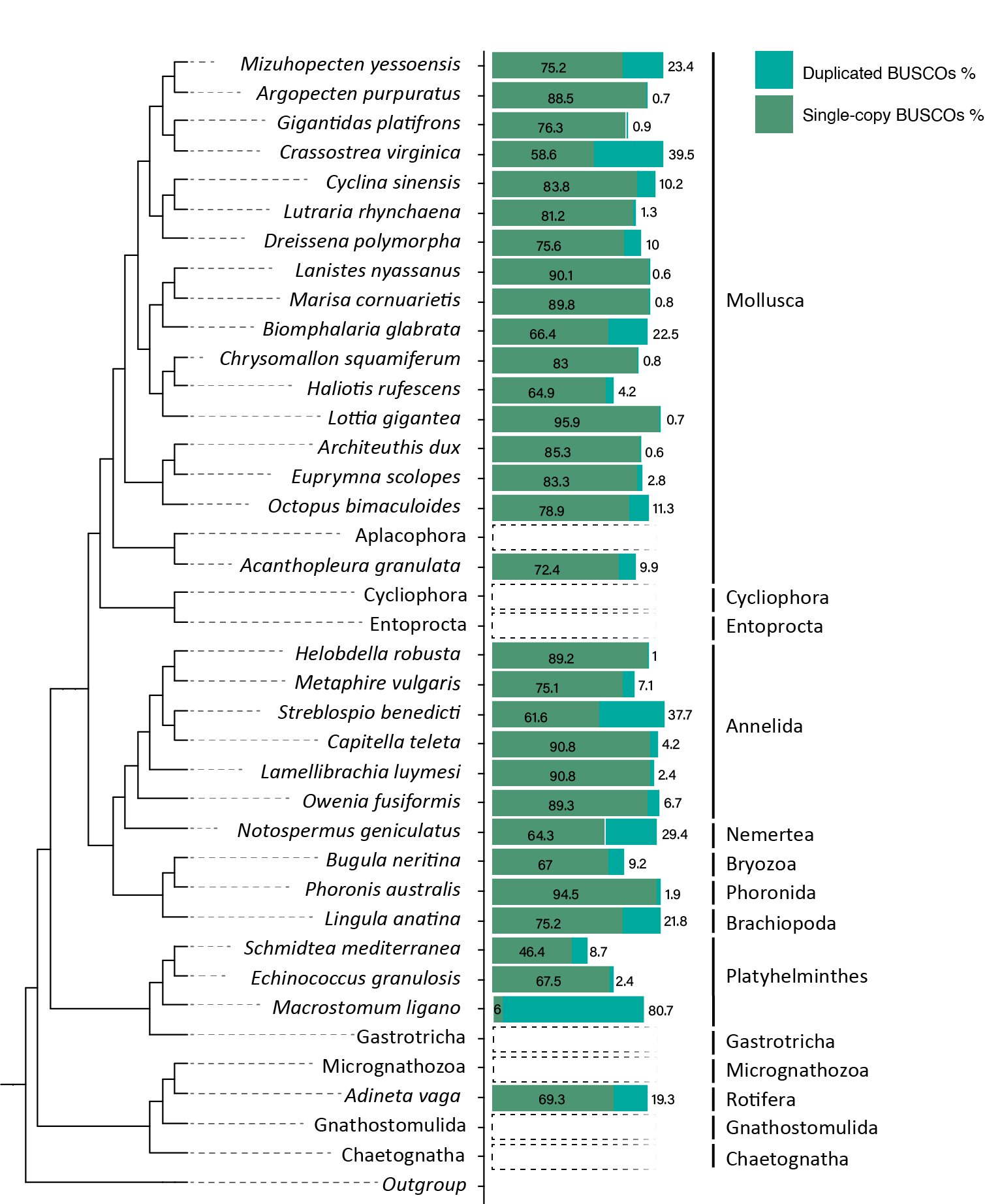

**Figure S1.** BUSCO scores of publicly available lophotrochozoan genomes (Metazoa odb10). Only genomes with a BUSCO completeness score >65% are included, except in the case of Platyhelminthes in which all genomes were included. Phyla with and without available data were placed in accordance with Laumer 2019.

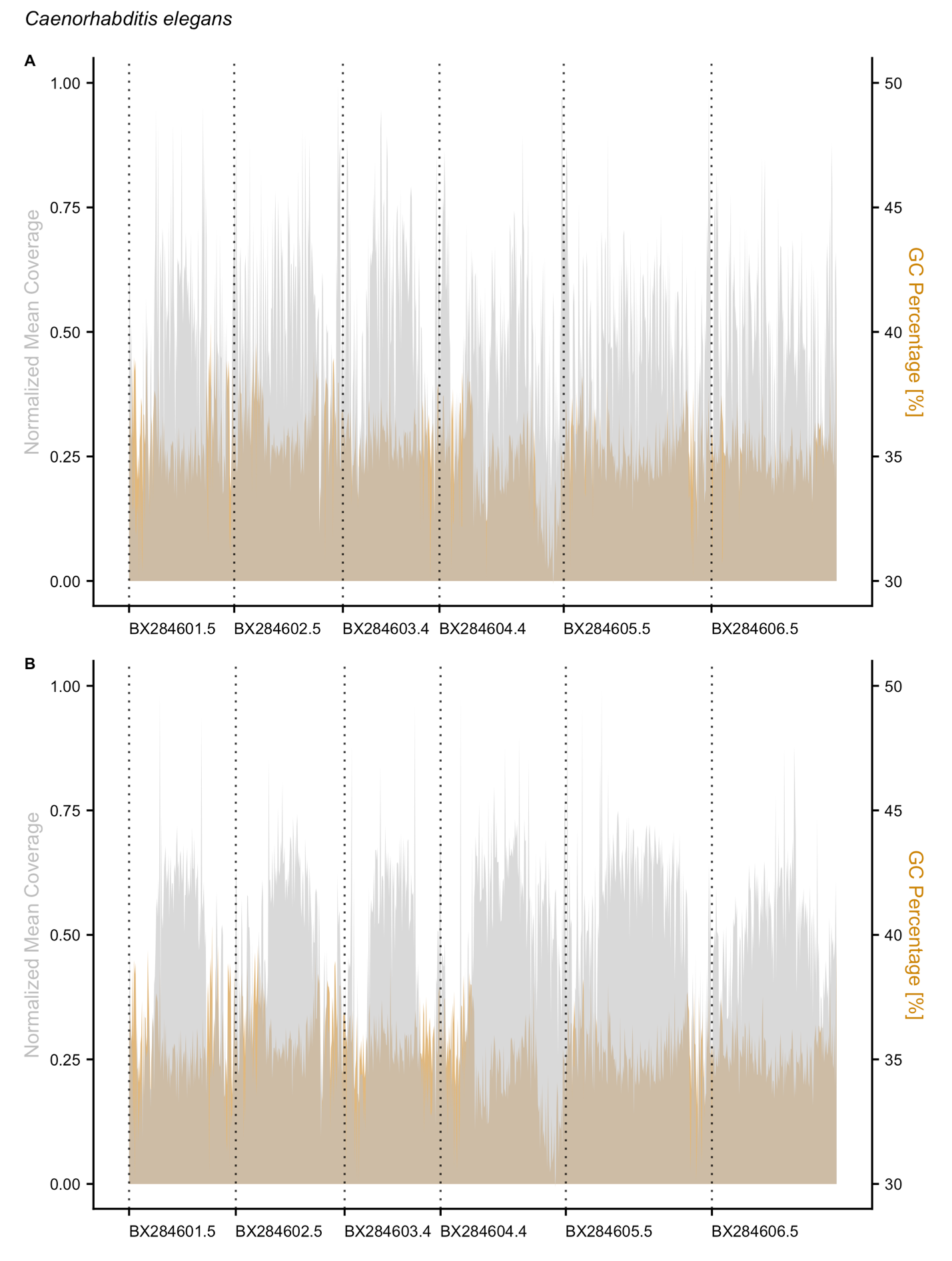

**Figure S2**: Mean coverage and GC content across 100 Kbp blocks of the *C. elegans* reference genome with outliers removed. A. Coverage of PacBio HiFi reads from MDA DNA aligned to the reference *C. elegans* genome [Gray] alongside GC percentage [Orange]. B. Coverage of unamplified PacBio HiFi reads aligned to the reference *C. elegans* genome [Gray] alongside GC percentage [Orange].

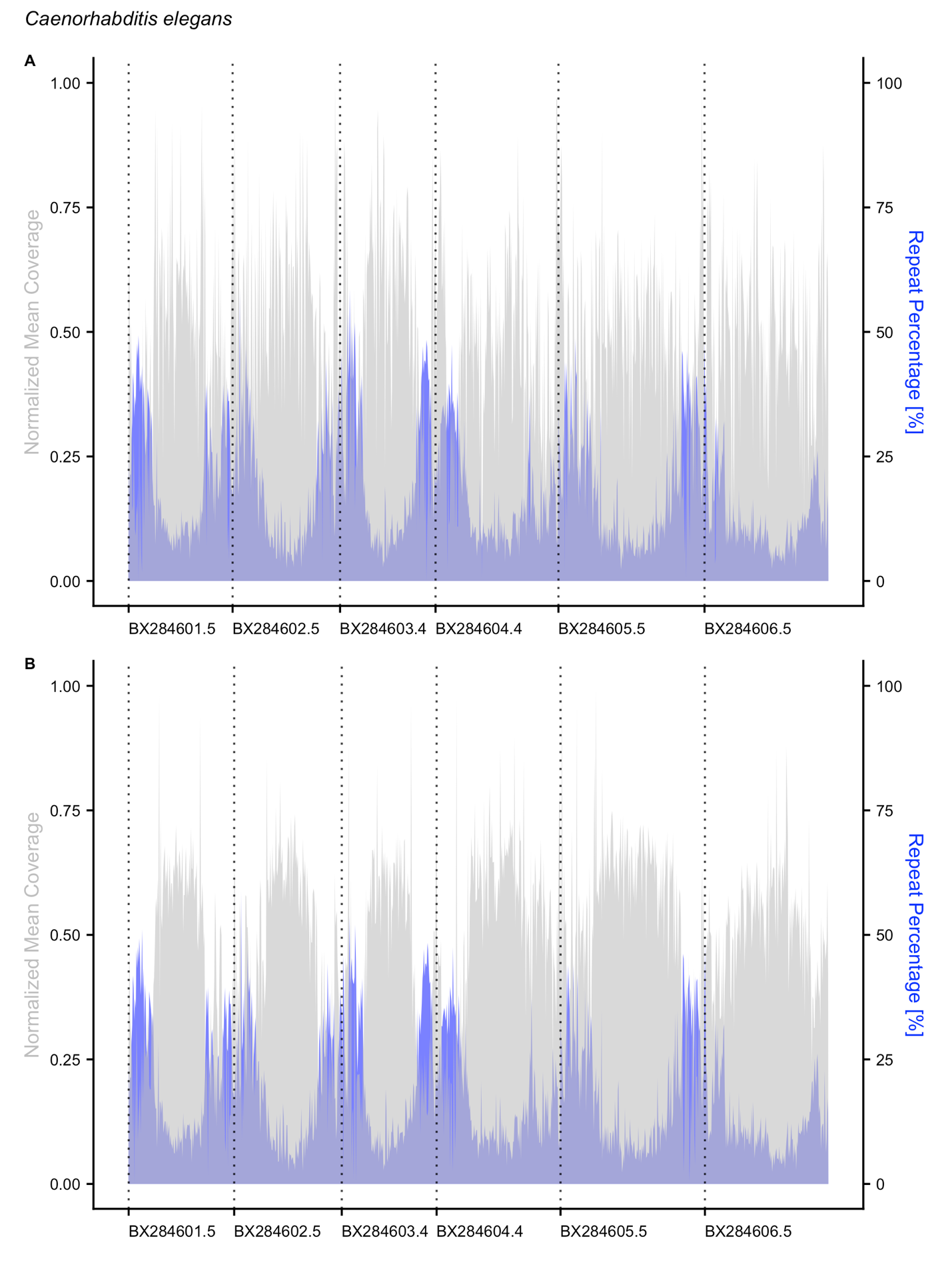

**Figure S3:** Mean coverage and repeat content across 100 Kbp blocks of the *C. elegans* reference genome with outliers removed. A. Coverage of PacBio HiFi reads from MDA DNA aligned to the reference *C. elegans* genome [Gray] alongside repeat content [Blue]. B. Coverage PacBio HiFi reads from unamplified DNA from a pool of worms aligned to the reference genome [Gray] alongside repeat content [Blue].

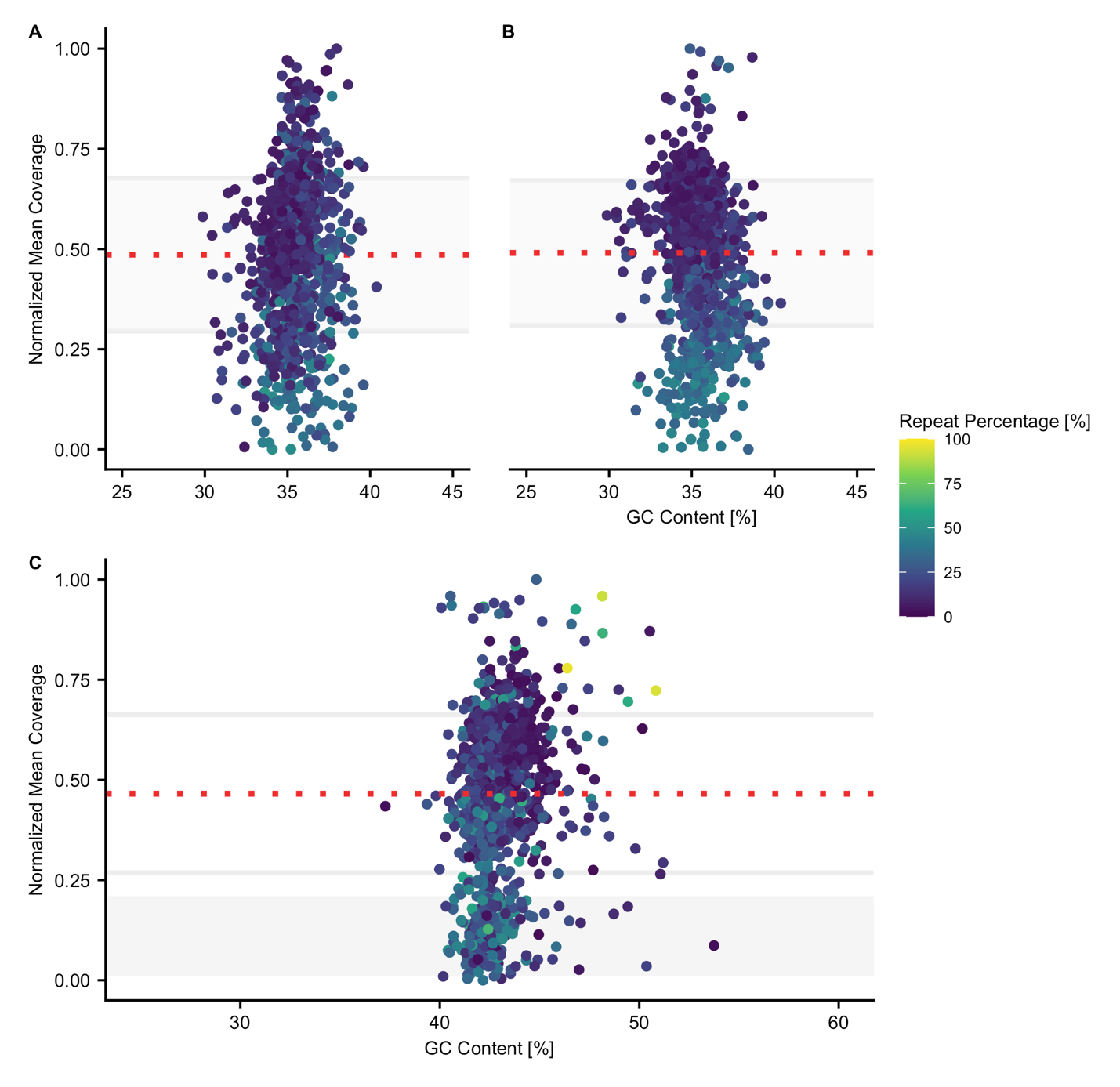

**Figure S4**: Normalized coverage, GC content, and repeat percentage of each genome assembly in 100kb blocks with outliers removed. A. Dotplot showing coverage of MDA amplified reads with respect to GC content for 100kb blocks of the *C. elegans* reference genome. B. Dotplot showing coverage of non-amplified reads from a pool of worms with respect to GC content for 100kb blocks of the *C. elegans* reference genome. C. Dotplot showing coverage with respect to GC content for contigs in the *L. squamata* genome assembly based on MDA DNA. A-C. Repeat content of each contig is indicated according to the key at the top right of the figure.

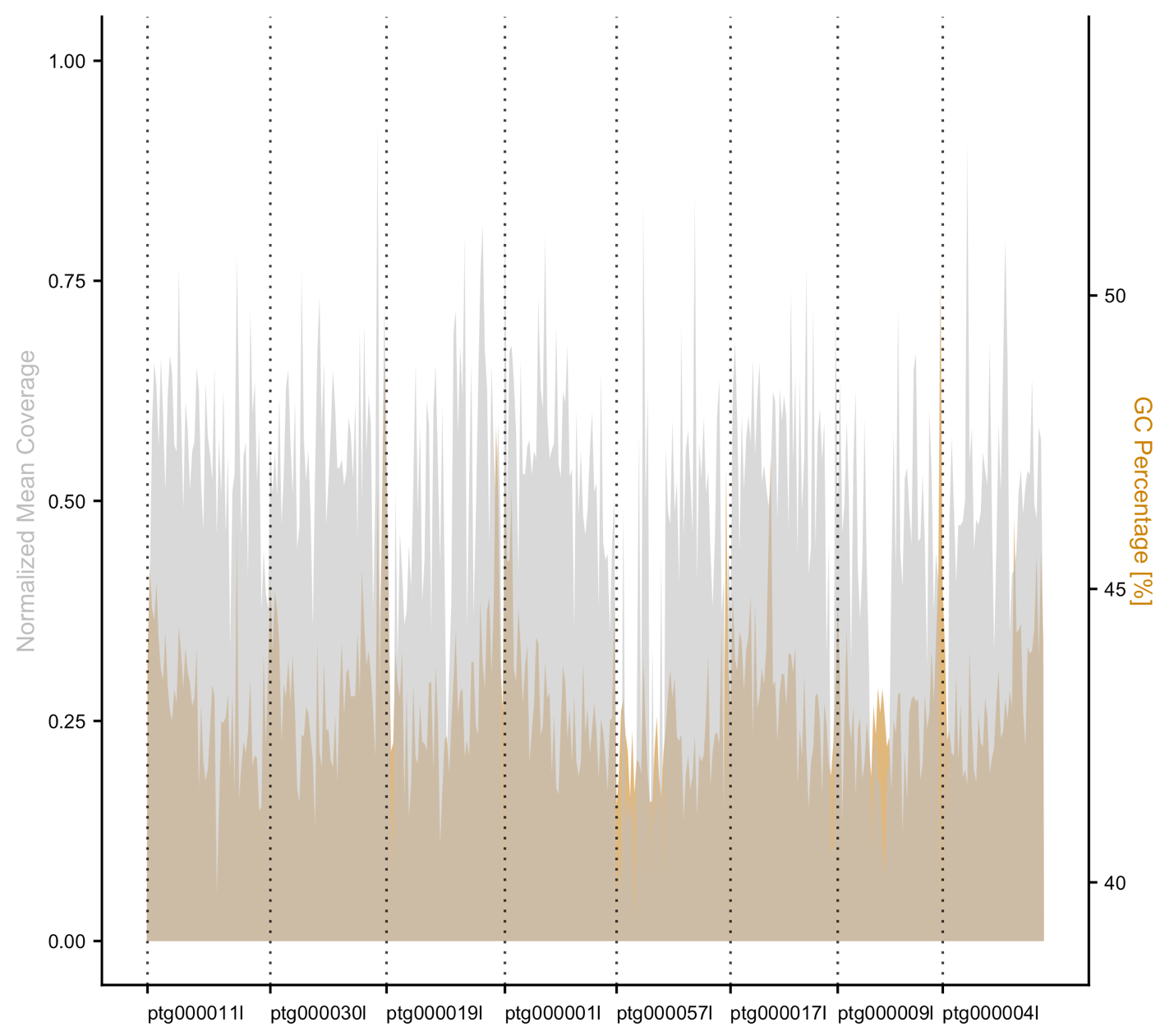

**Figure S5**: Coverage and GC content across 100kb blocks of the eight largest contigs of the

*L. squamata* genome with outliers removed. Coverage of PacBio HiFi reads from MDA DNA aligned to the 8 largest contigs of the *de novo* *L. squamata* genome assembly [Gray] alongside GC percentage [Orange].

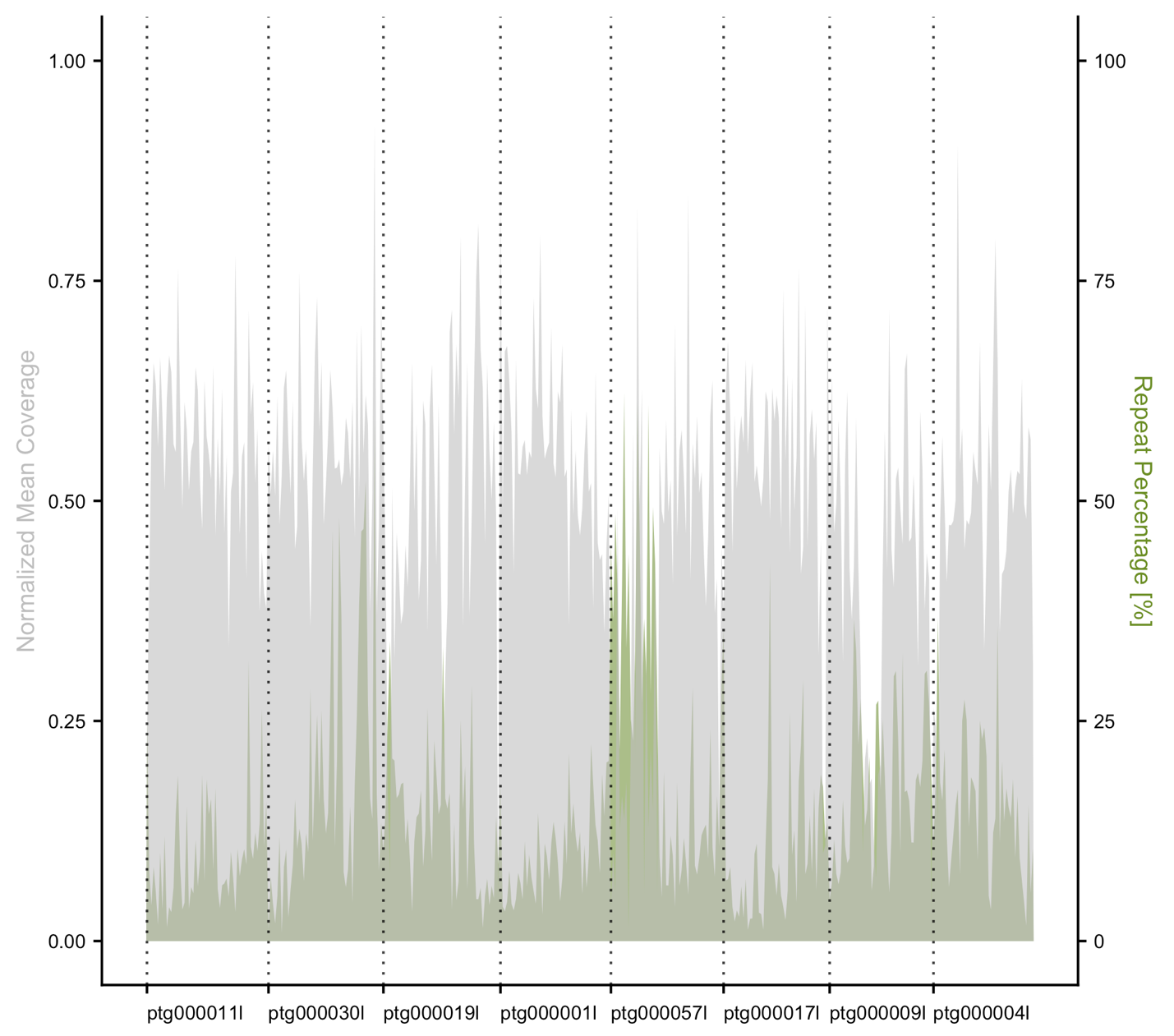

**Figure S6**: Coverage and repeat across 100 Kbp blocks of the eight largest contigs of the

*L. squamata* genome with outliers removed. Coverage of PacBio HiFi reads from MDA DNA aligned to the 8 largest contigs of the *de novo* *L. squamata* genome assembly [Gray] alongside repeat percentage [Green].
